# Hydrogel dimensionality directs mechanics-dependent nascent matrix deposition by tendon fibroblasts

**DOI:** 10.64898/2026.10.23.761444

**Authors:** Joshua Yu-Chung Liu, Ziqian Yang, Stephanie S. Steltzer, Mingyue Fan, Evelyn S. Navarro Salazar, Jiahao Liang, Megan L. Killian, Claudia Loebel

**Affiliations:** Department of Bioengineering, University of Pennsylvania, Philadelphia, USA; Center for Precision Engineering for Health (CPE4H), University of Pennsylvania, Philadelphia, USA; Department of Orthopaedic Surgery, University of Michigan, Ann Arbor, USA; Department of Molecular and Integrative Physiology, University of Michigan, Ann Arbor, USA

**Author notes:** Contributed equally (Joshua Yu-Chung Liu, Ziqian Yang).

## Abstract

Tendon fibroblasts continuously synthesize and remodel extracellular matrix (ECM) in response to their surrounding mechanical environment *in vivo* and *in vitro*, yet how dimensionality regulates the deposition of newly synthesized matrix remains poorly understood. Here, we investigated nascent matrix deposition by primary mouse tail tendon fibroblasts cultured on flat two-dimensional (2D) or encapsulated within three-dimensional (3D) norbornene-modified hyaluronic acid hydrogels with tunable mechanical properties. Newly synthesized matrix was visualized using metabolic labeling with azido-galactosamine (GalNAz), which labels nascent glycoproteins deposited during culture. In 2D, increasing hydrogel elastic modulus from 10 to 25 kPa enhanced nascent matrix deposition. This response was accompanied by increased cell spreading and YAP nuclear localization, indicating increased cellular mechanosensing. In contrast, tendon fibroblasts encapsulated within enzymatically degradable 3D hydrogels showed progressive matrix deposition over time but deposited less nascent matrix in stiffer (∼50 kPa) hydrogels compared with softer 5 and 10 kPa hydrogels. YAP nuclear localization was increased in softer hydrogels compared with 50 kPa hydrogels. Bulk RNA sequencing showed increased expression of tendon matrisome-associated genes in softer 3D hydrogels, including increased tenascin C and collagen type I expression. These findings demonstrate that hydrogel dimensionality determines the relationship between matrix mechanics and nascent matrix deposition. This work establishes a framework for studying how engineered microenvironments regulate the matrix-producing behavior of tendon fibroblasts and provides insight into the design of biomaterials for investigating tendon biology and matrix remodeling.

## Introduction

Tendons transmit forces between muscle and bone through a collagen-rich extracellular matrix (ECM)^1,2^. This matrix consists of type I collagen organized hierarchically into fibrils, fibers, and fascicles. It also contains collagen type III, proteoglycans and glycoproteins that regulate fibril assembly and tissue mechanics^3^. Tendon fibroblasts are the dominant cell type responsible for depositing and maintaining this ECM during homeostasis and injury repair^4,5^. Although the overall turnover of the mature collagenous matrix in adult tendon is relatively slow^6^, the matrix surrounding tendon cells and between fascicles is constantly synthesized and remodeled^7^. Tendon ECM deposition is therefore dynamic and responds to biological and environmental cues. In vivo, factors such as hypoxia^8,9^, mechanical tension^10,11^, and inflammation^12^ have been associated with altered tendon ECM during homeostasis and disease such as tendinopathy.

As cells within a mechanically active tissue, tendon fibroblasts continuously sense their mechanical environment through interactions with collagen fibers and the surrounding pericellular matrix^13,14^. Recent studies have linked changes in tendon loading to ECM remodeling.^15–18^ In particular, loss of mechanical tension in tissue has been shown to suppress YAP/TAZ signaling and increase catabolic activity^11^. In contrast, Rho/ROCK inhibition enhanced transforming growth factor beta 3 (TGF-β3)-induced tenogenesis of tendon progenitor cells and upregulation of tendon ECM genes^19^. However, it remains incompletely understood how the local mechanical microenvironment directs tendon fibroblast behavior and ECM deposition.

Hydrogel culture systems are a powerful approach for investigating how properties of the microenvironment regulate cell behavior^20,21^. Hydrogels are water-swollen polymer networks that are formed from natural ECM proteins or from engineered polymers such as hyaluronic acid^22,23^. Engineered hydrogels are useful as a reductionist platform because their modulus, cell adhesive moieties and degradability can be tuned^24–28^. This tunability has enabled studies of how hydrogel modulus regulates mechanotransduction, including cellular spreading and nuclear translocation of the transcriptional co-activator YAP^29^. Importantly, these responses depend on culture dimensionality. In 2D, increasing the modulus of the underlying hydrogel generally promotes cell spreading and YAP nuclear localization^30,31^. In 3D, however, increasing the hydrogel modulus is typically associated with greater mechanical confinement, which restricts cell spreading, and consequently reduces YAP nuclear localization^29^. Dimensionality therefore changes how cells sense and respond to matrix modulus.

In addition to responding to hydrogel properties, cells actively remodel their local microenvironment by depositing newly synthesized proteins at the cell-hydrogel interface, forming a layer of nascent ECM (nECM)^32,33^. Metabolic labeling with noncanonical amino acids or sugar analogs, and subsequent visualization with fluorophore-conjugated cyclooctynes via click chemistry, has enabled the characterization of this nECM^33,34^. Previous studies have shown that nECM deposition depends on hydrogel properties and that cells engage their own nECM in addition to the hydrogel^27,30–37^. However, it is unknown how hydrogel modulus and dimensionality together regulate nECM deposition.

Building upon these findings and the established relationship between mechanotransduction and tendon ECM remodeling, we used hydrogels to investigate how hydrogel modulus and dimensionality regulate tendon fibroblast nECM deposition. We used metabolic labeling to trace nECM from tendon fibroblasts in 2D and 3D hydrogel cultures with varying modulus. Our findings demonstrate that culture dimensionality determines the modulus-dependent mechanosensing response of tendon fibroblasts and shapes nECM deposition dynamics.

## Results and discussion

### Culture dimensionality determines how hydrogel moduli regulate tendon fibroblast spreading

Previous work showed that hydrogel modulus has opposing effects on mesenchymal stromal cells (MSCs) spreading and YAP nuclear localization in 2D and 3D hydrogels^29^. To determine whether tendon fibroblasts show similar dimensionality-dependent mechanosensing, we used the well-established and tunable norbornene-modified hyaluronic acid (NorHA) hydrogel system^43,44^. Tendon fibroblasts were isolated by outgrowth from the tail tendon explants of 6-8-week-old C57BL/6 male mice embedded in a collagen type I hydrogel (Fig. 1A). Cells were then seeded atop (2D) or encapsulated within (3D) NorHA hydrogels of varying modulus.

**Figure 1.**
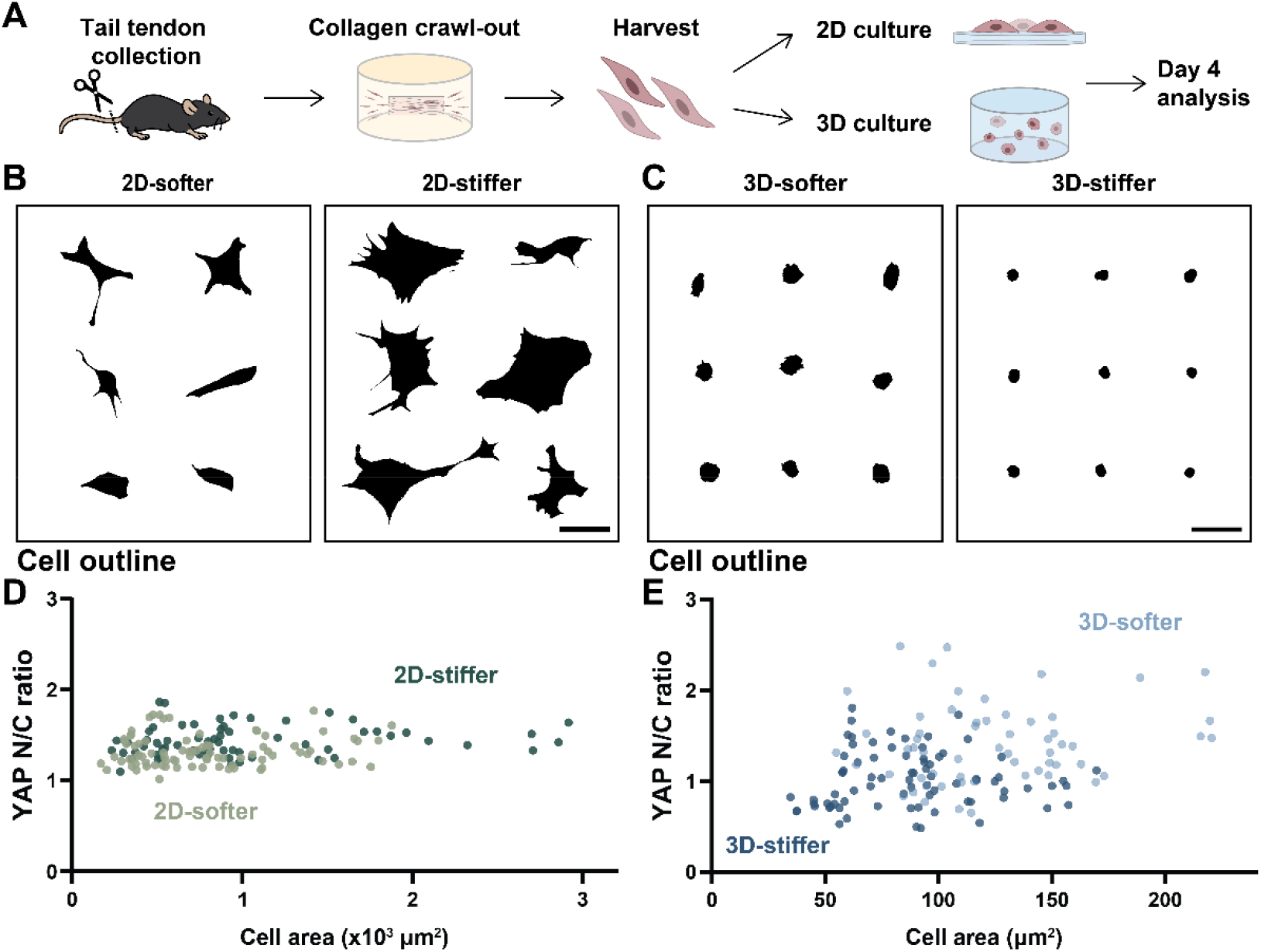
Hydrogel dimensionality alters tendon fibroblast morphology and mechanosensing. **A.** Schematic illustrating tendon fibroblast isolation using a collagen gel crawl-out method and subsequent culture in 2D and 3D hydrogel systems to investigate hydrogel modulus-dependent cellular responses. **B.** Representative binary masks showing cell outlines of tendon fibroblasts cultured on 2D-softer (10 kPa) and 2D-stiffer (25 kPa) hydrogels at day 4 (scale bar = 50 μm). **C.** Representative binary masks showing cell outlines of tendon fibroblasts cultured within 3D-softer (10 kPa) and 3D-stiffer (50 kPa) hydrogels at day 4 (scale bar = 50 μm). **D.** Scatter plot showing YAP nuclear-to-cytoplasmic (N/C) ratio as a function of cell area in 2D-softer (n = 84 cells, N = 2) and 2D-stiffer (n = 65 cells, N = 2) cultures. **E.** Scatter plot showing YAP N/C ratio as a function of cell area in 3D-softer (n = 59 cells, N = 2) and 3D-stiffer (n = 78 cells, N = 2) cultures. Dots represent individual cells. n = number of cells and N = number of independent experiments.

In 2D, tendon fibroblasts on stiffer hydrogels adopted more spread morphologies (Fig. 1B), which resulted in an increase in cell area on stiffer hydrogels compared to softer hydrogels (Fig. S1A). This result confirmed that cell area increases with hydrogel modulus in 2D culture. In contrast, cells cultured in 3D were generally smaller compared to 2D culture and stayed rounded throughout the 4 days of culture (Fig. 1C). Cell area of tendon fibroblasts within stiffer 3D hydrogels was decreased compared to those cultured in softer hydrogels (Fig. S1B), suggesting that increasing hydrogel modulus restricts cell spreading in 3D.

To investigate whether these morphological changes were associated with altered mechanosensing, we compared cell areas with YAP nuclear-to-cytoplasmic (N/C) ratio at the single-cell level. In 2D, cells cultured on the stiffer hydrogels spanned a broader range of cell areas than cells on the softer hydrogels, with substantial overlap in the distributions of YAP N/C ratios (Fig. 1D). In 3D, cells in softer hydrogels showed a broader distribution extending toward larger cell areas and higher YAP N/C ratios. In contrast, cells in stiffer hydrogels were more narrowly distributed toward smaller areas and lower YAP N/C ratios (Fig. 1E). Thus, increasing modulus promoted cell spreading in 2D, whereas softer hydrogels supported both spreading and YAP nuclear localization in 3D.

These findings indicate opposing modulus-dependent trends in tendon fibroblast spreading in 2D and 3D, with dimensionality-dependent differences in YAP localization, consistent with previous observations in MSCs^29^. This establishes the NorHA hydrogel system as a suitable platform for investigating how modulus and dimensionality regulate tendon fibroblast mechanosensing and nECM deposition.

### Hydrogel moduli regulate early YAP localization and nECM deposition in 2D

After having analyzed the relationship between cell area and YAP localization, we next quantified modulus-dependent differences in YAP localization over time and asked how these two parameters related to nECM deposition in 2D. NorHA was crosslinked with the non-degradable dithiol crosslinker dithiothreitol (DTT) to form mechanically tunable 2D hydrogels (Fig. 2A-i). Tendon fibroblasts at passage 2 were used throughout, as these cells deposited minimal nECM at passage 1 and showed signs of phenotypic changes such as a flatter morphology and enlarged cell area at passage 3. (Fig. S2). Cells were seeded at 4.4 × 10^3^ cells/cm^2^ density and cultured in 1% serum. nECM was metabolically labeled with the azide-modified sugar N-azidoacetylgalactosamine-tetraacylated (GalNAz, Fig. 2A-ii), which provided more detailed visualization of nECM than the methionine analog azidohomoalanine (AHA) (Fig. S3). Live-cell labeling with a DBCO-conjugated fluorophore then enabled visualization of newly deposited matrix surrounding individual cells (Fig. 2A-iii).

**Figure 2.**
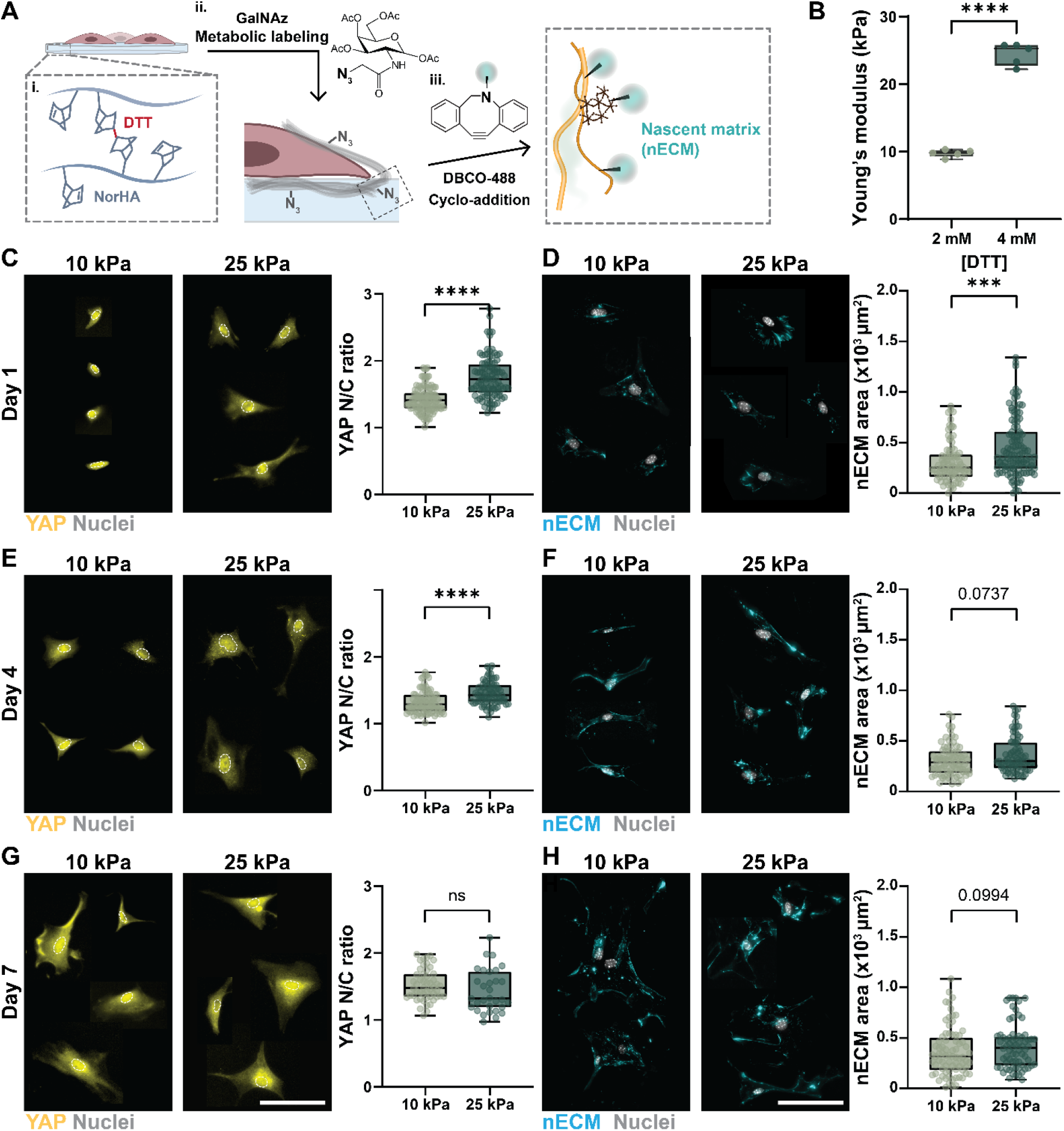
Higher 2D hydrogel modulus enhances YAP nuclear translocation and nECM deposition at early time points. **A.** Schematic illustrating the 2D norbornene-modified hyaluronic acid (NorHA) hydrogel system and metabolic labeling strategy used to visualize nascent extracellular matrix (nECM). NorHA hydrogels were crosslinked with dithiothreitol (DTT), and nECM was metabolically labeled using N-azidoacetylgalactosamine-tetraacylated (GalNAz), followed by dibenzocyclooctyne-488 (DBCO-488) cycloaddition for fluorescence visualization. **B.** Young’s modulus of hydrogels formed with 2 and 4 mM DTT crosslinker (*N* = 5 hydrogels per group). **C.** Representative images and quantification of YAP nuclear-to-cytoplasmic (N/C) ratio in tendon fibroblasts cultured on 10 and 25 kPa NorHA hydrogels at day 1 (10 kPa, *n* = 116 cells; 25 kPa, *n* = 101 cells; *N* = 2). **D.** Representative images and quantification of nECM area in tendon fibroblasts cultured on 10 and 25 kPa NorHA hydrogels at day 1 (10 kPa, *n* = 86 cells; 25 kPa, *n* = 113 cells; *N* = 2). **E.** Representative images and quantification of YAP N/C ratio in tendon fibroblasts cultured on 10 and 25 kPa NorHA hydrogels at day 4 (10 kPa, *n* = 87 cells; 25 kPa, *n* = 72 cells; *N* = 2). **F.** Representative images and quantification of nECM area in tendon fibroblasts cultured on 10 and 25 kPa hydrogels at day 4 (10 kPa, *n* = 78 cells; 25 kPa, *n* = 79 cells; *N* = 2). **G.** Representative images and quantification of YAP N/C ratio in tendon fibroblasts cultured on 10 and 25 kPa NorHA hydrogels at day 7 (10 kPa, *n* = 51 cells; 25 kPa, *n* = 31 cells; *N* = 2). **H.** Representative images and quantification of nECM area in tendon fibroblasts cultured on 10 and 25 kPa NorHA hydrogels at day 7 (10 kPa, *n* = 76 cells; 25 kPa, *n* = 70 cells; *N* = 2). Scale bar = 100 μm. Outliers were identified and excluded using the ROUT method (Q = 1%). *n* = number of cells and *N* = number of independent experiments. \*\*\*\**p* < 0.0001, \*\*\**p* < 0.001, ns = not significant, by unpaired Student’s *t*-test in B and Mann-Whitney test in C-H. Cell and nuclear boundaries used for YAP N/C quantification in C, E, and G are shown in Fig. S5.

NorHA was crosslinked with 2 or 4 mM DTT to fabricate hydrogels with elastic compression moduli (Young’s moduli) of 10 kPa (9.7 ± 0.5 kPa) and 25 kPa (24.5 ± 0.7 kPa, Fig. 2B). Within one day of culture, tendon fibroblasts attached with elongated morphologies on both hydrogels. However, cells on 25 kPa hydrogels showed greater spreading area and higher YAP N/C ratios than those on 10 kPa hydrogels (Fig. 2C). nECM deposition was already detectable at day 1 for both hydrogels with punctate structures below the cell body (Fig. S4). Quantification of total nECM area showed an approximately 1.46-folds increase for cells on 25 kPa compared to 10 kPa (Fig. 2D), suggesting that a higher YAP N/C ratio and increased cell spreading are associated with greater nECM deposition. By day 4, cells on both hydrogels became more elongated, and the difference in YAP N/C ratio between 10 and 25 kPa hydrogels decreased, although it remained significant (Fig 2E). The nECM formed more continuous and fibrous structures around the cells (Fig. S4), and the nECM area showed a trend towards higher values at 25 kPa (Fig 2F). By day 7, YAP N/C ratios were comparable between the two hydrogels (Fig. 2G). The nECM formed interconnected networks around cells on both hydrogels (Fig. S4), with only slightly higher nECM area for cells on 25 kPa (Fig 2H). Increased cell proliferation at this time point, however, limited our ability to distinguish nECM deposited by individual cells.

These results suggest that hydrogel modulus primarily regulates YAP localization and nECM deposition in 2D, whereas nECM organization progresses from punctate deposits to fibrous networks over time and independent of modulus.

### Culture conditions regulate tendon fibroblast viability and nECM deposition in 3D hydrogels

To extend our studies into 3D, NorHA was crosslinked with an MMP-cleavable peptide to allow cell-mediated degradation and spreading (Fig. 3A-i), and nECM was labeled with GalNAz and DBCO-fluorophore as in 2D (Fig. 3A-ii). Varying the crosslinker concentration (1, 2, or 7 mM) resulted in hydrogels with compression moduli of 5 kPa (4.4 ± 0.8 kPa), 10 kPa (10.6 ± 0.4 kPa), and 50 kPa (48.2 ± 5.6 kPa, Fig. 3B).

**Figure 3.**
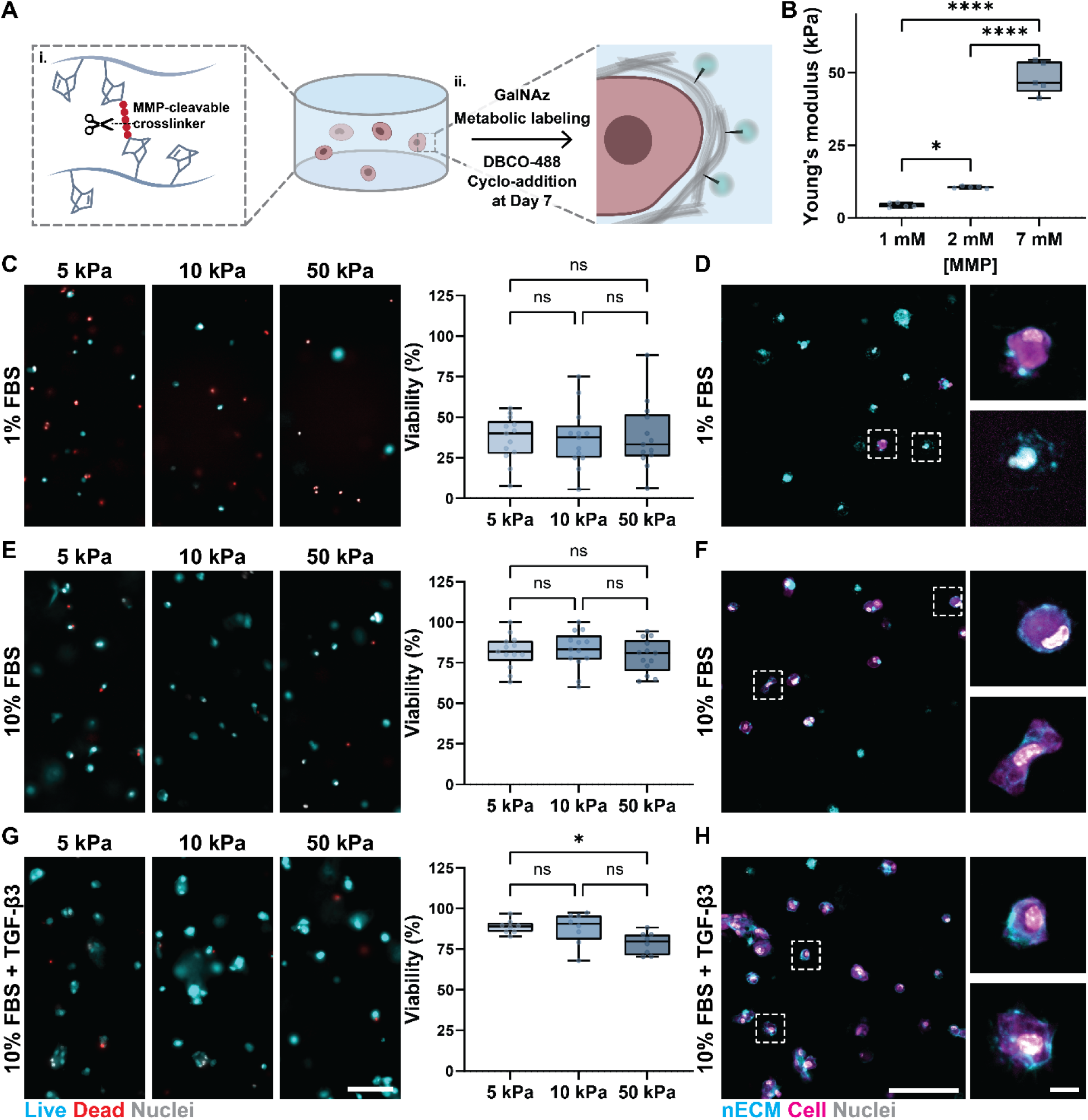
Enzymatically degradable NorHA hydrogels support 3D tendon fibroblast culture and nECM deposition. **A.** Schematic illustrating the 3D culture system consisting of NorHA hydrogels crosslinked with an MMP-cleavable dithiolated peptide crosslinker, followed by metabolic labeling and DBCO-mediated click chemistry to visualize nECM. **B.** Young’s modulus of hydrogels formed with 1, 2, and 7 mM MMP-cleavable crosslinker (N = 5 hydrogels per group). **C.** Representative images of Calcein AM (live cells), propidium iodide (dead cells), and Hoechst (nuclei) staining with quantification of tendon fibroblast viability after 7 days of culture in 5 kPa (1 mM), 10 kPa (2 mM), and 50 kPa (7 mM) NorHA hydrogels supplemented with 1% FBS (n = 13 fields of view per group, N = 3 independent experiments). **D.** Representative images of nECM deposition by tendon fibroblasts cultured for 7 days in 10 kPa NorHA hydrogels supplemented with 1% FBS. **E.** Representative images and quantification of tendon fibroblast viability after 7 days of culture in 5, 10, and 50 kPa NorHA hydrogels supplemented with 10% FBS (n = 13 fields of view per group, N = 3 independent experiments). **F.** Representative images of nECM deposition by tendon fibroblasts cultured for 7 days in 10 kPa NorHA hydrogels supplemented with 10% FBS. **G.** Representative images and quantification of tendon fibroblast viability after 7 days of culture in 5, 10, and 50 kPa hydrogels supplemented with 10% FBS and TGF-β3 (n = 8 fields of view per group, N = 2 independent experiments). **H.** Representative images of nECM deposition by tendon fibroblasts cultured for 7 days in 10 kPa NorHA hydrogels supplemented with 10% FBS and TGF-β3. Dashed boxes in D, F, and H indicate cells shown at higher magnification to the right (viability images, scale bar = 100 μm; nECM main images, scale bar = 100 μm; magnified images, scale bar = 10 μm). n = number of fields of view and N = number of independent experiments. ***p < 0.0001, p < 0.05, ns = not significant by one-way ANOVA with Tukey’s multiple comparisons test.

We first encapsulated tendon fibroblasts in 3D hydrogels followed by culture in media containing 1% FBS as used for 2D culture, but cell viability was below 40% after 7 days of culture and across all hydrogels (Fig. 3C). Most cells showed collapsed morphologies, although a small population of viable cells desposited detectable nECM (Fig. 3D). Increasing the serum concentration to 10% FBS improved cell viability to approximately 80% across all hydrogels (Fig. 3E). Most cells deposited nECM around the cell body by day 7 (Fig. 3F). As transforming growth factor beta-3 (TGF-β3) has been reported to promote tenogenic differentiation^19^, we further supplementated the media containing 10% FBS with 50 ng/mL TGF-β3^45^. Viability remained high, although it was reduced in 50 kPa hydrogels compared to 5 kPa hydrogels (Fig. 3G). The majority of cells deposited a pericellular nECM layer (Fig. 3H). However, TGF-β3 also increased cell clustering, which could introduce cell-cell interactions that confound the effects of cell-hydrogel interactions. We therefore used 10% FBS without TGF-β3 for subsequent 3D experiments.

### Hydrogel moduli shape the extent and timing of nECM deposition in 3D

We next asked how hydrogel modulus regulates nECM deposition over time in 3D. In 5 kPa hydrogels, nECM was discontinuous and concentrated in discrete regions around the cells at day 1 and formed a more continuous pericellular layer with fibrous structures extending into the surrounding hydrogel at day 4 and day 7 (Fig. 4A). Quantification of nECM area showed an increase from day 1 to day 4 with little difference at day 7 (Fig. 4B).

**Figure 4.**
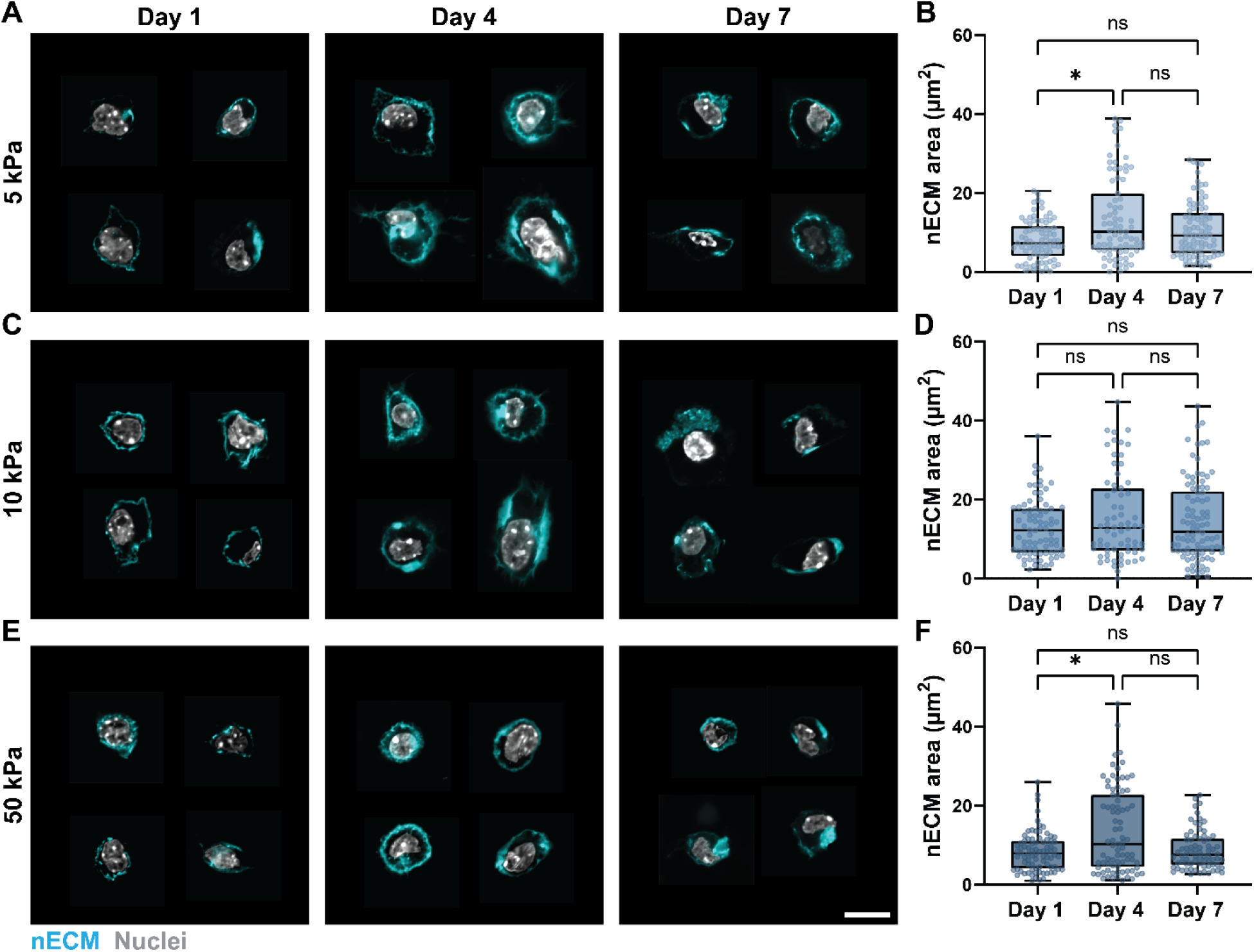
nECM deposition shows modulus-dependent temporal dynamics in 3D hydrogels. **A.** Representative images of nECM deposition by tendon fibroblasts in 5 kPa NorHA hydrogels over 7 days of culture. **B.** Quantification of nECM area in 5 kPa hydrogels (day 1 n = 81 cells, N = 2; day 4 n = 79 cells, N = 2; day 7 n = 89 cells, N = 2). **C.** Representative images of nECM deposition in 10 kPa NorHA hydrogels. **D.** Quantification of nECM area in 10 kPa hydrogels (day 1 n = 81 cells, N = 2; day 4 n = 68 cells, N = 2; day 7 n = 94 cells, N = 2). **E.** Representative images of nECM deposition in 50 kPa NorHA hydrogels. **F.** Quantification of nECM area in 50 kPa hydrogels (day 1 n = 82 cells, N = 2; day 4 n = 81 cells, N = 2; day 7 n = 74 cells, N = 2). Scale bar = 10 μm. n = number of cells and N = number of independent experiments. *p < 0.05, ns = not significant by Kruskal–Wallis test with Dunn’s multiple comparisons test.

In 10 kPa, a continuous nECM was already apparent at day 1, with larger nECM regions and fibrous structures extending into the surrounding hydrogel at day 4 and day 7 (Fig. 4C). Quantification of nECM area showed no detectable change over time (Fig. 4D). In 50 kPa, nECM was sparse and confined close to the cell body at day 1 and formed a more continuous pericellular layer with fewer fibrous structures at day 4 and day 7 (Fig. 4E). Similar to 5 kPa hydrogels, nECM area in 50 kPa hydrogels increased from day 1 to day 4 with little changes at day 7 (Fig. 4F).

Previous studies have shown greater nECM area in softer hydrogels, likely due to increased diffusion of nECM into the crosslinked network^41^. Here, nECM was higher in 10 kPa compared to 5 kPa hydrogels, suggesting that additional cellular responses contribute to modulus-dependent nECM deposition. In addition, the decrease in nECM area from day 4 to day 7 in 5 and 50 kPa hydrogels may reflect diffusion of nECM beyond the detectable pericellular region or cell-mediated remodeling and degradation of nECM within the MMP-degradable hydrogels. Future studies will further need to determine whether nECM deposition changes more gradually at moduli between 10 and 50 kPa.

### Hydrogel moduli regulate matrisome gene expression in 3D

Beyond total nECM deposition amount, we next asked whether hydrogel modulus alters the ECM transcriptional program of tendon fibroblasts using bulk RNA sequencing of cells cultured in 3D hydrogels for 4 days. Principal component analysis (PCA) separated the 50 kPa condition from the largely overlapping 5 and 10 kPa conditions (Fig. 5A). MatrisomeDB classification^46,47^ identified 554 matrisome genes, including 165 core matrisome and 389 matrisome-associated genes (Fig. 5B). Consistent with the PCA, only 17 genes were differentially expressed (DEGs; |log2FC| > 1, adjusted *p* < 0.1) between 5 and 10 kPa hydrogels (Fig. 5C). In contrast, 382 and 334 DEGs were identified when comparing 50 kPa to 5 and 10 kPa hydrogels, with more matrisome genes enriched within the 5 kPa hydrogels (51 enriched at 5 kPa vs. 26 at 50 kPa, Fig. 5D) compared to 10 kPa hydrogels (42 enriched at 10 kPa vs. 16 at 50 kPa, Fig. 5E).

**Figure 5.**
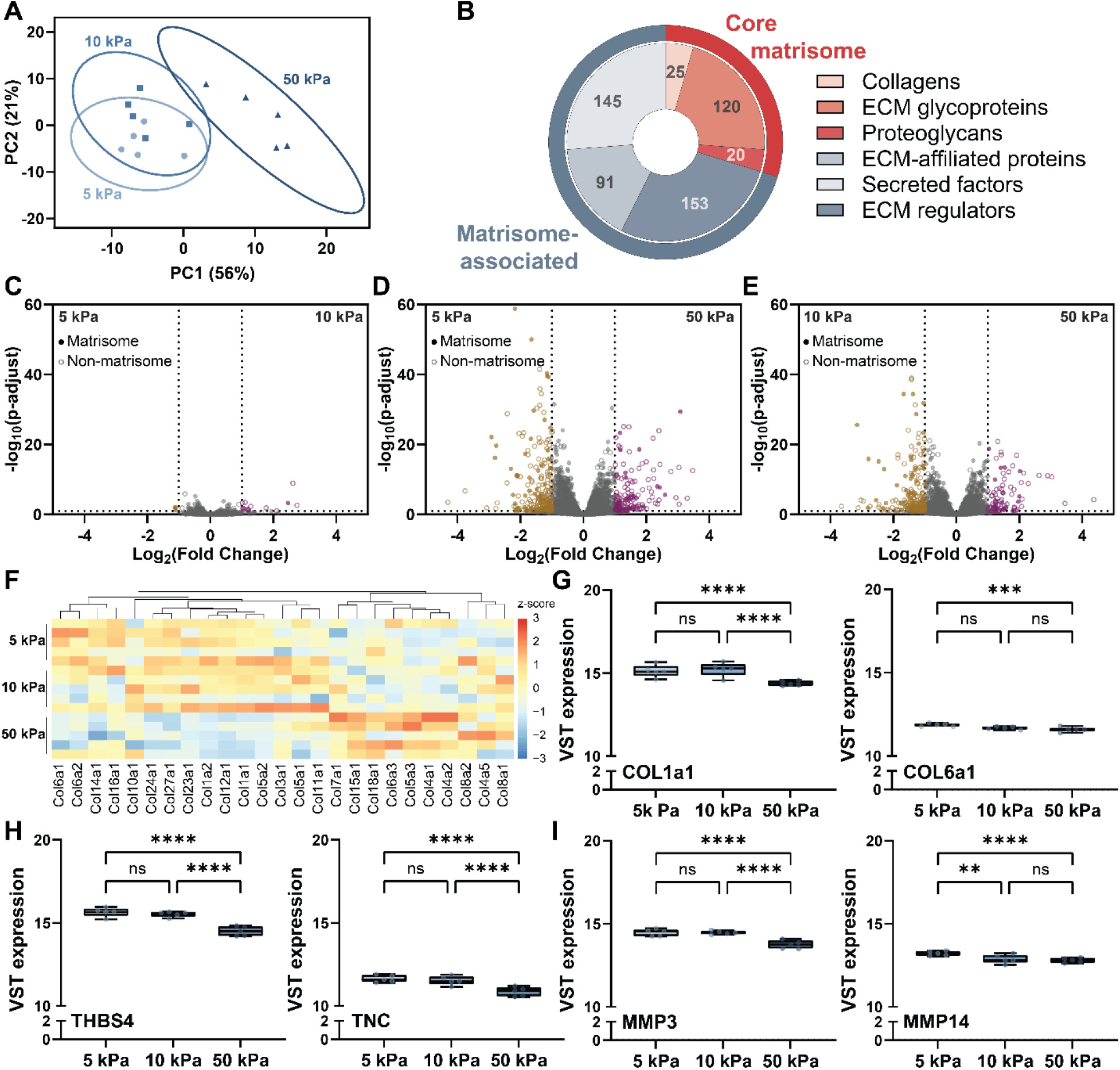
Lower 3D hydrogel modulus promotes expression of tendon-associated matrisome genes. **A.** Principal component analysis (PCA) of transcriptomic profiles of tendon fibroblasts cultured in 1, 2, and 7 mM MMP-crosslinked NorHA hydrogels (5, 10, and 50 kPa, respectively) at day 4 (*N* = 5 independent samples per group). Ellipses indicate 95% confidence regions. **B.** Distribution of identified matrisome genes, including core matrisome components (collagens, ECM glycoproteins, and proteoglycans) and matrisome-associated components (ECM-affiliated proteins, secreted factors, and ECM regulators). **C-E.** Volcano plots showing differential gene expression between **(C)** 5 and 10 kPa, **(D)** 5 and 50 kPa, and **(E)** 10 and 50 kPa NorHA hydrogels. Filled circles indicate matrisome genes and open circles indicate non-matrisome genes. Yellow indicates genes enriched in the lower modulus condition and violet indicates genes enriched in the higher modulus condition; gray indicates genes not meeting the significance threshold. Differentially expressed genes were defined as adjusted *p* < 0.1 and |log2 fold change| > 1. **F.** Heatmap showing VST-transformed expression of collagen genes across 5, 10, and 50 kPa hydrogels. Expression values were standardized by gene (row Z-score), with genes ordered by hierarchical clustering. **G.** VST-transformed expression of *Col1a1* and *Col6a1*. **H.** VST-transformed expression of tendon-associated ECM genes *Thbs4* and *Tnc*. **I.** VST-transformed expression of matrix catabolic genes *Mmp3* and *Mmp14*. A, C-I. *N* = 5 independent samples per group. **G-I**, \*\*\*\**p* < 0.0001, \*\*\**p* < 0.001, \*\**p* < 0.01, ns = not significant by DESeq2 pairwise differential-expression analysis with Benjamini-Hochberg-adjusted *p* values (FDR).

Within the collagen family, fibrillar collagens such as type I were generally enriched at 5 and 10 kPa, whereas several non-fibrillar collagens, including types IV and VIII showed higher expression at 50 kPa (Fig. 5F). Modulus-dependent enrichment was also evident across other ECM categories, although proteoglycans showed a less pronounced stiffness-dependent pattern. (Fig. S6). Among tendon-associated ECM genes, *Col1a1* was reduced at 50 kPa compared to 5 and 10 kPa, and *Col6a1*, a major component of the tendon fibroblast pericellular matrix^13^ was reduced at 50 kPa compared to 5 kPa (Fig. 5G). *Col3a1* was relatively consistent across moduli, whereas the basement membrane collagen Col4a1 increased at 50 kPa (Fig S7.). The expression of tendon-enriched matricellular protein *Thbs4* and the glycoprotein *Tnc*, were also reduced at 50 kPa (Fig. 5H). In addition, expression of the matrix metalloproteinases *Mmp3* and *Mmp14* decreased with increasing modulus (Fig. 5I).

Because nECM area increased between 5 and 10 kPa hydrogels with minimal transcriptional changes, hydrogel properties likely affect the assembly or retention of nECM. In contrast, the 50 kPa hydrogels induced a distinct transcriptional state with reduced expression of fibrillar and tendon-associated ECM genes that may contribute to lower nECM area at high modulus.

### Hydrogel moduli alter YAP and tendon-associated ECM localization

To determine whether this transcriptional shift was associated with altered mechanosensing, we next measured YAP localization in 3D hydrogels at day 4. YAP nuclear localization was similar between 5 and 10 kPa hydrogels but reduced in 50 kPa hydrogels (Fig. 6A), indicating greater YAP activation in softer 3D microenvironments.

**Figure 6.**
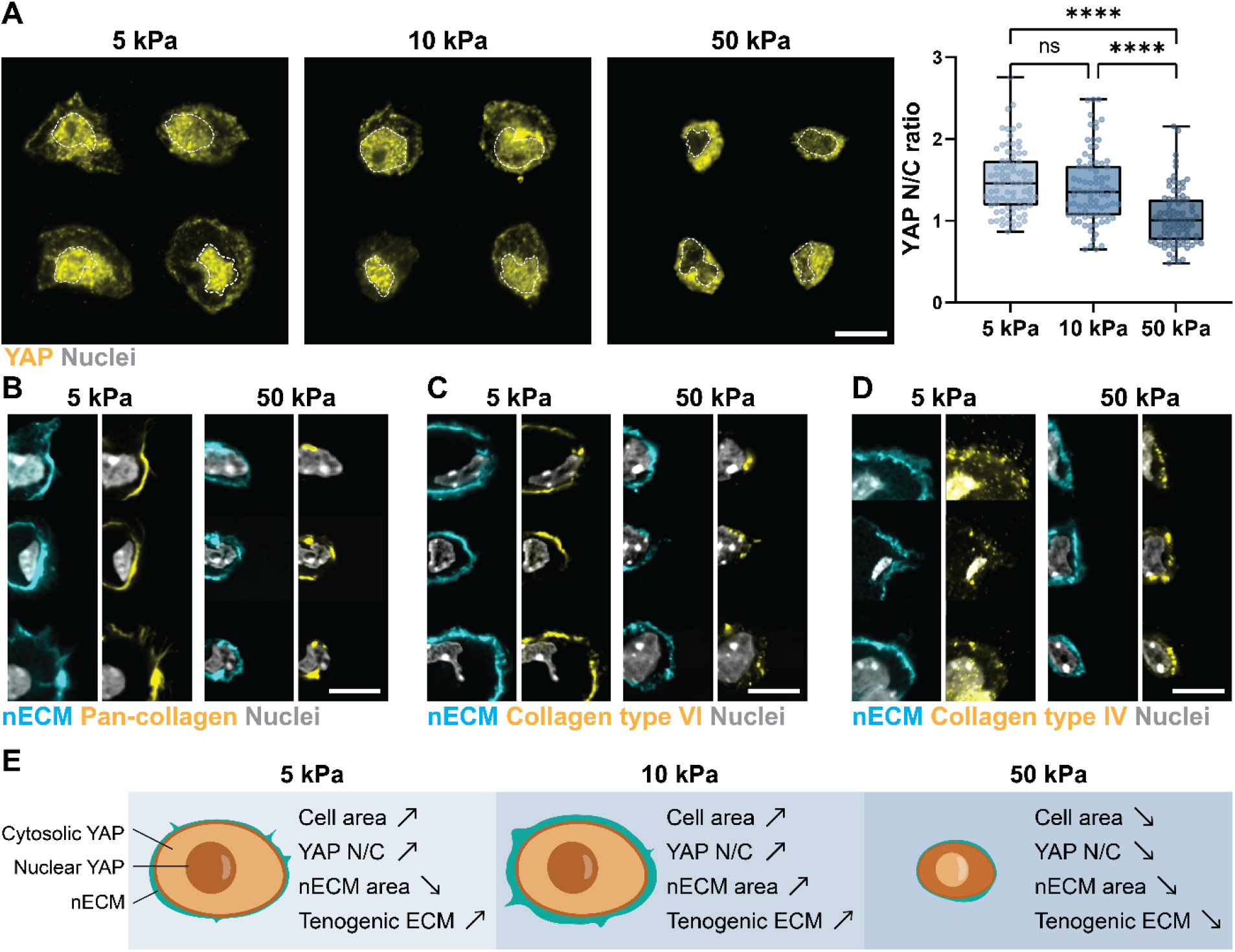
Lower 3D hydrogel modulus supports YAP nuclear translocation and deposition of tendon-associated pericellular matrix. **A.** Representative images and quantification of YAP N/C ratio in tendon fibroblasts cultured within 5, 10, and 50 kPa NorHA hydrogels (5 kPa, *n* = 83 cells; 10 kPa, *n* = 82 cells; 50 kPa, *n* = 82 cells; *N* = 2). White dashed line indicates the outline of the nucleus. Representative images of **B.** pan-collagen, **C.** collagen type VI and **D.** collagen type IV with nECM and nuclei of tendon fibroblasts cultured within 5 kPa and 50 kPa hydrogels. **E.** Schematic summarizing modulus-dependent changes in cell area, YAP N/C, nECM and tenogenic ECM expression in 5, 10, and 50 kPa 3D NorHA hydrogels. \*\*\*\**p* < 0.0001; ns = not significant.

As transcript abundance does not always reflect ECM protein deposition^7,48^, we next investigated the spatial organization of specific ECM components by immunostaining. Since 5 and 10 kPa hydrogels showed similar YAP localization and transcriptional profiles, we focused this analysis on 5 and 50 kPa hydrogels. Pan-collagen staining largely overlapped with nECM labeling in both conditions, but formed continuous, fiber-like structures in 5 kPa hydrogels and more punctate deposits in 50 kPa hydrogels (Fig. 6B). Similarly, collagen type VI, a major component of the tendon pericellular matrix, formed a continuous pericellular layer in 5 kPa hydrogels and more punctate structures with less continuous coverage in 50 kPa hydrogels (Fig. 6C). In contrast, the basement membrane collagen type IV was predominantly diffuse in 5 kPa hydrogels but concentrated within the pericellular nECM in 50 kPa hydrogels (Fig. 6D). Thus, reduced YAP nuclear localization at high modulus in 3D was associated with a shift from a continuous collagen type VI-rich pericellular matrix to collagen type IV deposition. These findings suggest that 3D confinement redirects tendon fibroblasts away from a tendon-associated matrix. In summary, these results show that 5 and 10 kPa hydrogels support cell spreading, YAP nuclear localization, and a tendon-associated ECM profile compared to 50 kPa hydrogels. Notably, nECM area was the greatest at the intermediate 10 kPa modulus, indicating that nECM deposition is not determined by transcriptional changes alone (Fig. 6E).

### Conclusion and outlook

In this study, we used NorHA hydrogels and metabolic labeling to determine how hydrogel modulus and culture dimensionality regulate nECM deposition by tendon fibroblasts. Our results showed in 2D, higher substrate modulus transiently enhanced YAP nuclear localization and nECM deposition, and these differences diminished as nECM matured into fibrous networks. In 3D, the relationship between modulus and nECM was less linear. Cells in 5 and 10 kPa hydrogels showed similar YAP localization and transcriptional profiles, yet nECM area was greatest and sustained over time at 10 kPa, suggesting that hydrogel properties affecting nECM assembly or retention may also regulate nECM accumulation. In contrast, fibroblasts in 50 kPa hydrogels had reduced YAP nuclear localization with decreased expression of tenogenic ECM-affiliated genes and less deposition of tendon-associated, collagen type VI-rich pericellular matrix. Interestingly, *Mmp3* and *Mmp14* expression also decreased with increasing modulus. Previous studies of tendon detensioning showed increased ECM catabolism with elevated *Mmp3* expression, and this was also associated with softer tendons^11^. Our results suggest that reduced YAP activity from 3D confinement, similar to what has been reported with reduced tension in explanted tissue, may regulate matrix remodeling through distinct mechanisms. Since our hydrogels are MMP-degradable, lower MMP expression in cells seeded in 50 kPa hydrogels may further limit cell-mediated hydrogel degradation and spreading, reinforcing the confined, low-YAP state and the reduced nECM area at high modulus. Future studies should explore how independently tuning modulus and degradability^49,50^ of hydrogels contributes to distinct cell-intrinsic responses from nECM retention.

Direct comparison across dimensionalities is challenging, as 2D and 3D cultures often require different serum concentrations to maintain cell viability^51,52^. The diffusion gradients of 3D culture that may limit serum availability may also limit oxygen availability, which has known effects on nECM deposition of tendon fibroblasts^8^. Additional studies are required to probe the effects of serum and oxygen concentrations on the interplay between tendon fibroblast proliferation, differentiation and nECM deposition^53^. It is also important to note that cells were isolated from tail tendons of young mice and cultured under static conditions, which do not capture the mechanical loading or regional heterogeneity of tendons *in vivo*. Several recent advances in dynamic 3D tendon culture systems such as dynamic loading of *ex vivo* tendon explants^54^ or microtissues^55^ may be beneficial to test the influence of different mechanical modes. Finally, combining metabolic labeling with nascent proteomics^33^ could further identify which ECM components are deposited under each condition.

## Experimental

### Primary tendon fibroblasts isolation and culture

Primary tendon fibroblasts were isolated from tail tendons of 6-8-week-old male C57BL/6 mice obtained from The Jackson Laboratory. All animal procedures were approved by the University of Pennsylvania Institutional Animal Care and Use Committee (IACUC; protocol no. 807632). For each independent isolation, tail tendons from three mice were pooled.

Tendon explants were embedded in 3 mg/mL bovine type I collagen gels (FibriCol®, Advanced BioMatrix) and cultured for 10 days to allow tendon fibroblasts to migrate and proliferate from the tissue into the surrounding collagen hydrogel. Collagen gels were then digested with 0.25 mg/mL collagenase type II (Worthington Biochemical) for 40 min at 37 °C to recover the migrated cells. Cells recovered from the collagen gels were defined as passage 1 (P1) and expanded in Dulbecco’s Modified Eagle Medium/F12 (Gibco) supplemented with 10% fetal bovine serum (FBS, Corning) on standard tissue culture plastic (Thermo Fisher Scientific). Unless otherwise indicated, P2 cells were used for all experiments, and cells were not cryopreserved.

### Synthesis of norbornene-modified hyaluronic acid

Norbornene-modified hyaluronic acid (NorHA) was synthesized as previously described^44^. Briefly, sodium hyaluronate (MW ∼ 68.1kDa, Lifecore Biomedical) was dissolved at 1% (w/v) in 2-(N-morpholino)ethanesulfonic acid (MES, Sigma-Aldrich) buffer (pH 5.5). 4-(4,6-Dimethoxy-1,3,5-triazin-2-yl)-4-methylmorpholinium chloride (DMTMM, TCI) and 5-norbornene-2-methylamine (Nor, TCI) were sequentially added at a 1:3:2 molar ratio of HA repeat units:DMTMM:Nor, and the reaction was stirred for 24 h at room temperature. Saturated sodium chloride (Sigma-Aldrich) solution was then added at 8% (v/v), and the modified polymer was precipitated in ethanol and collected by vacuum filtration. The polymer was redissolved in deionized water, dialyzed against deionized water for 72 h using dialysis tubing with a molecular weight cutoff of 6-8 kDa, and lyophilized. The degree of norbornene modification was determined by proton nuclear magnetic resonance (^1H NMR) spectroscopy using a 600 MHz NEO spectrometer (Bruker) and was approximately 23%, as quantified using MestReNova (v15.1.0, Mestrelab Research).

### 2D NorHA hydrogel fabrication and tendon fibroblast seeding

NorHA hydrogels were fabricated on coverglass for 2D tendon fibroblast culture. To covalently attach the hydrogels, 12-mm circular coverglasses were methacrylated for 6 min at room temperature in ethanol containing 0.3% (v/v) glacial acetic acid (Thermo Fisher Scientific) and 0.5% (v/v) 3-(trimethoxysilyl)propyl methacrylate (MilliporeSigma) and air-dried for 30 min. Hydrogel precursor solutions contained 4 wt% NorHA, 2 or 4 mM dithiothreitol (DTT, Sigma-Aldrich), 1 mM thiolate RGD peptide (GCGYGRGDSPG; GenScript), and 1.7 mM lithium phenyl-2,4,6-trimethylbenzoylphosphinate (LAP, Arkema Sartomer). Precursor solutions were pipetted onto Sigmacote(Sigma-Aldrich)-treated coverglass, overlaid with methacrylated 12-mm circular coverglasses, and photocrosslinked with UV light (365 nm, 5 mW/cm²) for 3 min. Hydrogel-coated coverglasses were removed and swollen overnight in phosphate-buffered saline (PBS) before cell seeding. For glass controls, 12-mm glass coverglasses were plasma-treated and coated with 100 µg/mL type I collagen in 0.01 M HCl. Tendon fibroblasts were seeded at 5 × 10³ cells per coverglass. For metabolic labeling with GalNAz, cells were cultured in DMEM/F12 supplemented with 1% FBS, 50 µM tetra acetylated N-azidoacetylgalactosamine (GalNAz, Vector Laboratories), and 50 µg/mL ascorbate-2-phosphate (MilliporeSigma). For metabolic labeling with AHA, cells were cultured in a 1:1 mixture of F-12 and methionine-free DMEM (Gibco) supplemented with 1% FBS, 25 µM L-azidohomoalanine (AHA, Vector Laboratories), 75 µM L-methionine (Sigma-Aldrich), and 50 µg/mL ascorbate-2-phosphate. Culture media was changed every other day.

### 3D NorHA hydrogel fabrication and tendon fibroblast encapsulation

NorHA hydrogels were fabricated for 3D culture as previously reported^33^. Lyophilized NorHA was sterilized by germicidal UV exposure for 30 min. Hydrogel precursor solutions contained 4 wt% NorHA, 1 mM thiolated RGD peptide, 1.7 mM LAP, and 1, 2, or 7 mM MMP-degradable dithiol peptide crosslinker (GCNS*VPMSMRGG*SNCG; GenScript). Tendon fibroblasts were suspended in precursor solution at 5 × 10^⁶^ cells/mL and photocrosslinked with UV light (365 nm, 5 mW/cm²) for 3 min. Cell-laden hydrogels were cultured in DMEM/F12 supplemented with 1% FBS, 10% FBS, or 10% FBS with 50 ng/mL transforming growth factor beta-3 (TGFβ-3, R&D system). All media contained 50 µM GalNAz and 50 µg/mL ascorbate-2-phosphate for metabolic labeling of nECM. Culture media was changed every other day.

### Cell viability

Cell viability in 3D hydrogels was assessed at day 4 by incubating samples with 2 µM calcein AM (Invitrogen), 5 µg/mL propidium iodide (PI; MilliporeSigma), and Hoechst 33342 (1:1000, Thermo Fisher Scientific) in PBS for 30 min at 37°C. Samples were washed with PBS and imaged as z-stacks on a Leica THUNDER microscope with a 25× objective.

### Click labeling and immunofluorescence staining

Azide-labeled nECM was visualized by live-cell click labeling with 30 µM AZDye 488 DBCO (Vector Laboratories) in PBS containing 2% bovine serum albumin (BSA, MilliporeSigma) at 37°C. Samples were then washed with PBS and fixed in 4% paraformaldehyde (PFA, Thermo Fisher Scientific) for 30 min at room temperature.

Fixed 3D hydrogels were incubated overnight in 30% sucrose, infiltrated with optimal cutting temperature (OCT, Thermo Fisher Scientific) for 2 h at 4°C, embedded in OCT, and snap-frozen in 2-methylbutane(TCI) cooled with liquid nitrogen. Hydrogels were cryosectioned at 10 µm (CM3050S, Leica), and sections were baked for 30 min at 37°C and washed with PBS to remove residual OCT.

For immunostaining, hydrogels were permeabilized with 0.1% Triton X-100 for 10 min (YAP staining only), blocked with 5% BSA for 30 min, and incubated overnight at 4°C with primary antibodies against YAP (Santa Cruz Biotechnology, sc-101199; 1:100), anti-collagen IV (Invitrogen, PA1-85320; 1:200), or anti-collagen VI (Biosynth, 70R-CR009X; 1:200) in 2% BSA. Samples were then incubated for 1 h at room temperature with goat anti-mouse or goat anti-rabbit IgG Alexa Fluor 568 secondary antibodies (A11031 and A11011, Invitrogen; 1:200), together with HCS CellMask Deep Red (H32721, Invitrogen; 1:5000) where indicated. For pan-collagen, 3D sections were heated in PBS at 95 °C for 25 min to denature collagen^56^. Cy3-conjugated collagen hybridizing peptide (R-CHP, 3Helix, 20 µM in PBS) was heated at 80 °C for 5 min, rapidly cooled on ice, and applied to the sections, and incubated overnight at 4 °C.

3D sections were mounted in SlowFade Diamond Antifade Mountant with DAPI (Invitrogen), and 2D samples were imaged in PBS. 2D samples were imaged on a Leica THUNDER microscope with a 25× objective, and 3D sections on a Nikon Eclipse Ti2-E microscope with a 60× objective. Identical acquisition settings were used across conditions.

### Image analysis

All image analyses were performed in ImageJ (v1.54p, NIH). Cell viability was calculated as the percentage of Hoechst-positive cells that were PI-negative. For cell area, whole-cell masks were generated from the HCS CellMask Deep Red channel. For YAP quantification, nuclear masks were generated from the DAPI or Hoechst channel, and cytoplasmic regions were defined by subtracting the nuclear mask from the whole-cell mask. The YAP N/C ratio was calculated as the mean YAP nuclear intensity divided by the mean cytoplasmic YAP intensity. For nECM quantification, background in the DBCO channel was subtracted using a rolling-ball radius of 50 pixels, and DBCO-positive regions were segmented using Otsu thresholding. Objects smaller than 5 pixels² were excluded, and nECM area was measured for each cell.

### RNA isolation, sequencing, and analysis

At day 4, cell-laden hydrogels were snap-frozen and mechanically minced in TRIzol reagent (Thermo Fisher Scientific) on ice. Following chloroform (Thermo Fisher Scientific) extraction to remove hydrogel debris, total RNA was purified using the Direct-zol RNA Miniprep kit with on-column DNase digestion and the RNA Clean & Concentrator kit (both Zymo Research) according to the manufacturer’s instructions. RNA concentration and purity were assessed using a NanoDrop spectrophotometer (Thermo Fisher Scientific), and RNA integrity was assessed using a TapeStation (Agilent), with RINe values of 9.6-10.0 for all samples. RNA was shipped in SEQguard Dino Preserve (Genovis) in TE buffer (pH 7.5) to Plasmidsaurus for 3′ end-counting library preparation and sequencing.

Reads were quality-filtered using fastp (v0.24.0), aligned to the mouse reference genome (GRCm39; Ensembl; GCA_000001635.9) using STAR (v2.7.11b), deduplicated based on unique molecular identifiers (UMIs) using UMIcollapse (v1.1.0), and quantified at the gene level using featureCounts from the Subread package (v2.1.1). Differential expression analysis was performed using DESeq2 (v1.42.1) in R (v4.3.2). Analyses were restricted to protein-coding genes, and genes with fewer than 10 counts in at least two samples were excluded. Variance-stabilizing transformation (VST) was used for principal component analysis and heatmaps. Differential expressions between 5, 10, and 50 kPa hydrogels were assessed using pairwise analysis. Differentially expressed genes were defined as those with Benjamini-Hochberg-adjusted *p* value < 0.1 and |log_2_(fold change)| > 1. Matrisome genes were annotated using the mouse MatrisomeDB database^46,47^.

### Statistical analysis

Statistical analyses were performed in GraphPad Prism (version 11.1.0) and R (version 2026.08.2). Outliers were removed using the ROUT method (Q = 1%), and normality was assessed using the Shapiro-Wilk test. Two groups were compared using unpaired two-tailed Student’s t-tests or Mann-Whitney tests for normally and non-normally distributed data. Three or more groups were compared using one-way ANOVA with Tukey’s post hoc test or Kruskal-Wallis tests with Dunn’s post hoc test, as appropriate.

### Author Contributions

J.Y. Liu and Z. Yang developed the study design, conducted the experiments, performed data analysis and visualization, and wrote the initial manuscript. S. S. Steltzer contributed to the development and establishment of the tenocyte isolation and culture protocol and provided scientific input. M. Fan, E. S. Navarro Salazar and J. Liang assisted with experimental studies and data collection. M.L. Killian provided scientific input and contributed to the interpretation of the findings. C. Loebel directed and supervised the study, contributed to its conceptual development and interpretation, and provided critical revisions to the manuscript. M.L. Killian and C. Loebel acquired funding supporting this work. All authors reviewed and edited the manuscript and approved the final version.

### Generative AI Disclosure Statement

Generative AI (ChatGPT, GPT-5.6 Sol, OpenAI) was used solely to refine the language, including grammar, wording, and sentence clarity. All suggested edits were reviewed by the authors to ensure the accuracy of the final text.

## Acknowledgements

This work was partially supported by the National Institutes of Health (R00-HL151670 and R35GM157063 to C.L.; R01AR082348 to M.L.K.), a pilot award through Penn PAT-CORT (P50AR080581 to C.L.) and the David and Lucile Packard Foundation (to C.L.).

## Data Availability Statement

The RNA sequencing data generated in this study have been deposited in the Dryad Digital Repository under https://doi.org/10.5061/dryad.t4b8gtjk6. Additional data supporting the findings of this study are available from the corresponding author upon reasonable request.

