## Supplementary Figures for "Hydrogel dimensionality directs mechanics-dependent nascent matrix deposition by tendon fibroblasts"

^#^Contributed equally

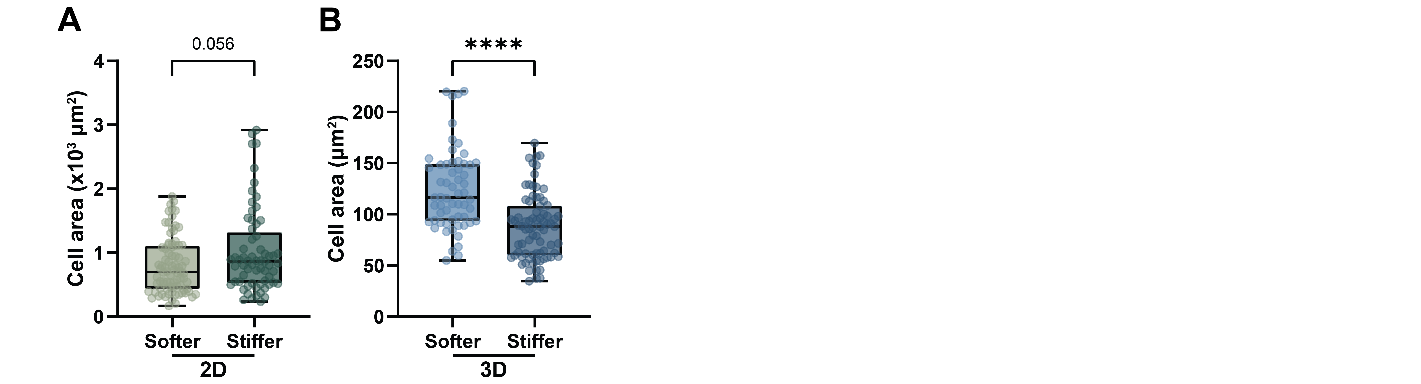

**Supplementary Figure 1. Cell area of tendon fibroblasts in 2D and 3D NorHA hydrogels at day 4.** **A.** Quantification of cell area of tendon fibroblasts cultured on softer (10 kPa) and stiffer (25 kPa) NorHA hydrogels at day 4 (softer, *n* = 84 cells; stiffer, *n* = 65 cells; *N* = 2). **B.** Quantification of cell area of tendon fibroblasts cultured within softer (10 kPa) and stiffer (50 kPa) NorHA hydrogels at day 4 (softer, *n* = 59 cells; stiffer, *n* = 78 cells; *N* = 2). *n* = number of cells and *N* = number of independent experiments. ****p* < 0.0001, **p* < 0.05 by Welch’s *t*-test

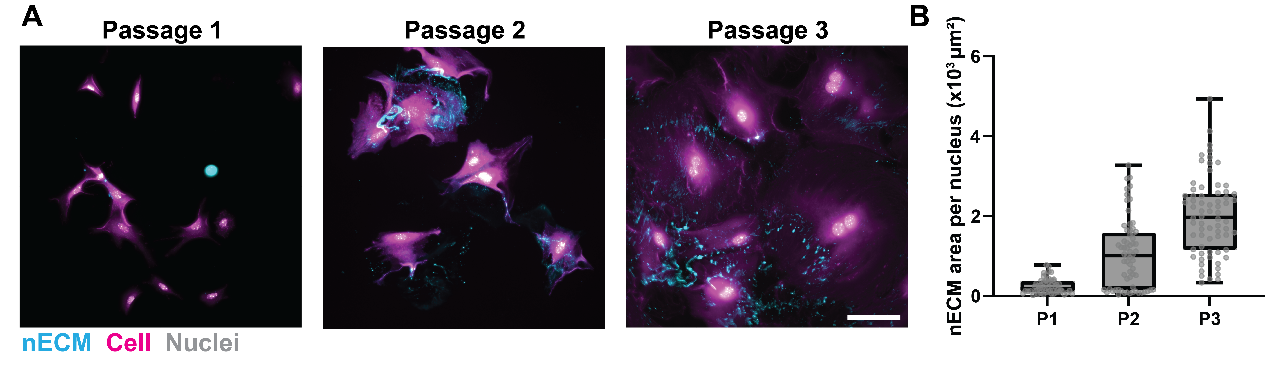

**Supplementary Figure 2. Metabolic labeling of nECM in tendon fibroblasts across passages. A**. Representative images of nECM, cell body, and nuclei in passage 1-3 tendon fibroblasts cultured on glass for 4 days in medium containing tetraacetylated N-azidoacetylgalactosamine (GalNAz), followed by DBCO-488 cycloaddition. Scale bar = 100 μm. **B**. Quantification of nECM area normalized to the number of nuclei per field of view (FOV; P1, n = 53 FOVs; P2, n = 65 FOVs; P3, n = 70 FOVs; N = 3 independent experiments). P1-P3 indicate passages 1-3. *****p* < 0.0001 by Kruskal-Wallis test followed by Dunn’s multiple-comparisons test.

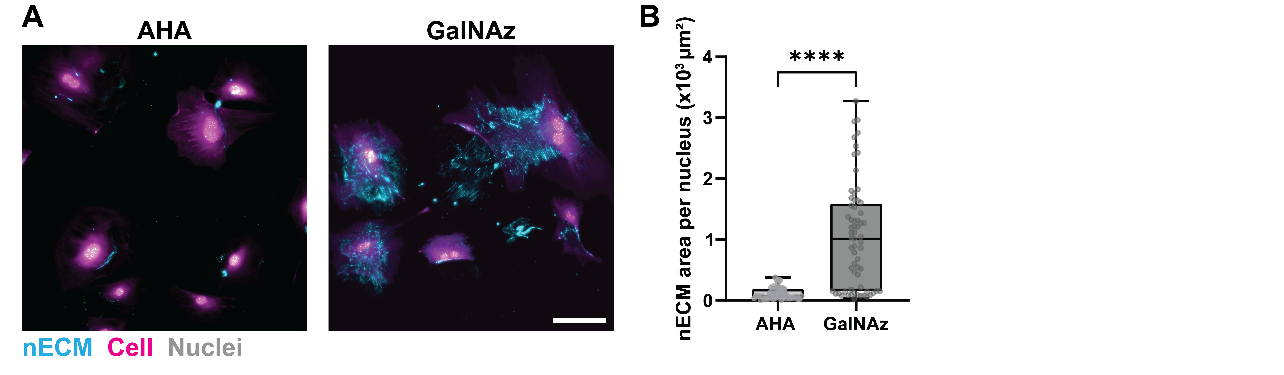

**Supplementary Figure 3. Metabolic labeling of nECM in tendon fibroblasts using AHA and GalNAz. A.** Representative images of nECM in tendon fibroblasts cultured on glass for 4 days in medium containing azidohomoalanine (AHA) and tetraacetylated N-azidoacetylgalactosamine (GalNAz), followed by DBCO-fluorophore cycloaddition. Scale bar = 100 μm. **B.** Quantification of nECM area normalized to the number of nuceli per field of view (FOV, AHA, n = 54 FOVs; GalNAz, n = 65 FOVs; N = 3 independent experiments). *****p* < 0.0001 by two-tailed Mann-Whitney test.

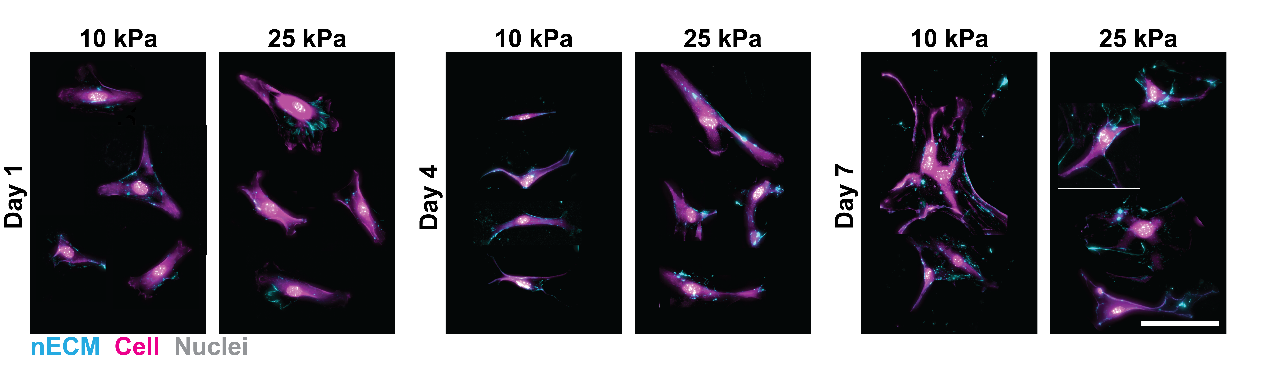

**Supplementary Figure 4. Co-staining of nECM and cell body in 2D tendon fibroblast culture.** Representative images of nECM, cell body, and nuclei in tendon fibroblasts cultured on 10 and 25 kPa NorHA hydrogels for 1, 4, and 7 days. Scale bar = 100 μm.

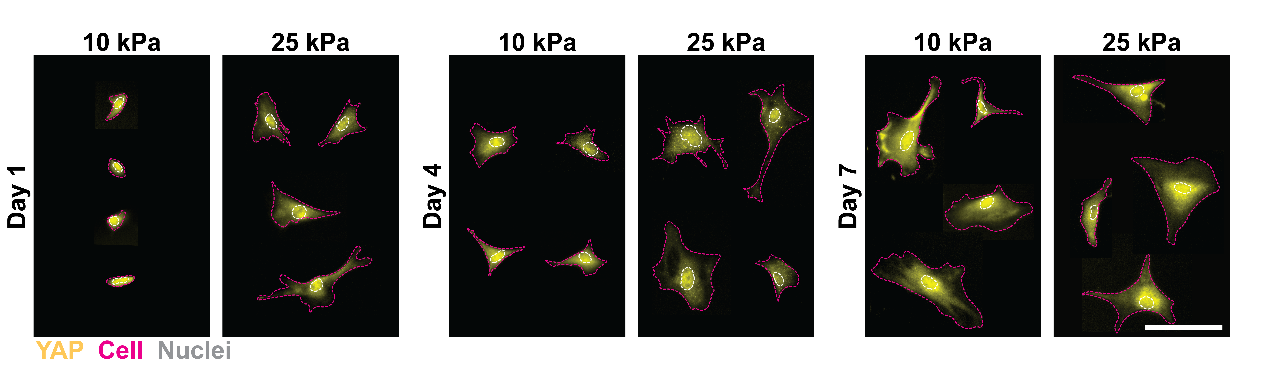

**Supplementary Figure 5. YAP staining with cell and nuclear outlines in 2D tendon fibroblast culture.** Representative images of YAP immunostaining in tendon fibroblasts cultured on 10 and 25 kPa NorHA hydrogels for 1, 4, and 7 days. Cell and nuclear boundaries used for YAP N/C quantification are indicated by magenta dashed and white outlines. Scale bar = 100 μm

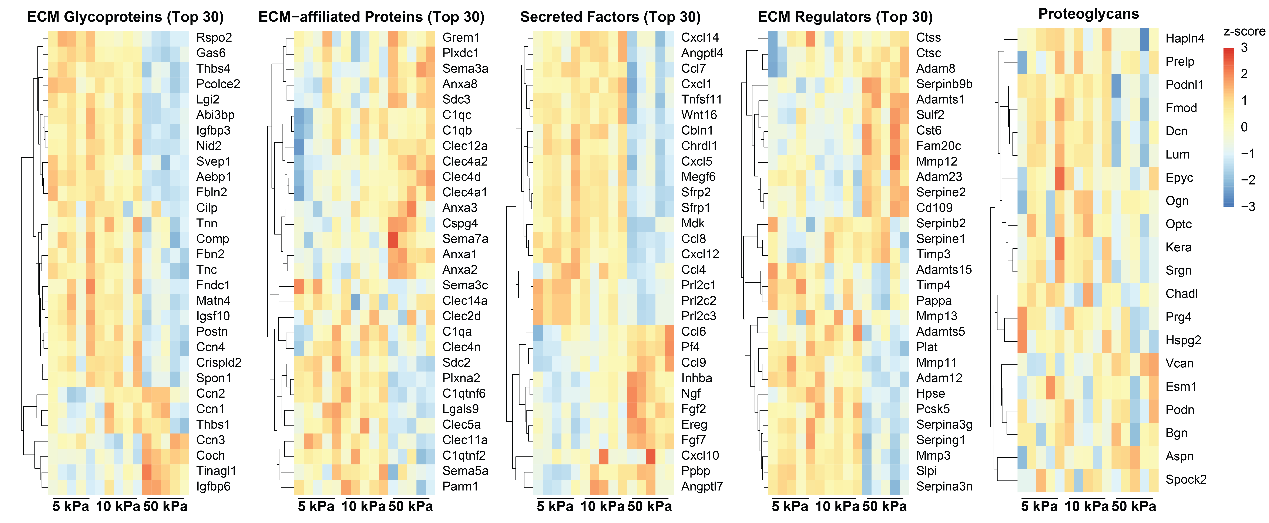

**Supplementary Figure 6. Heatmaps of gene expression in 3D tendon fibroblast culture.** Heatmaps of the 30 most variable genes in each of the ECM glycoproteins, ECM-affiliated proteins, secreted factors, and ECM regulators, and of all identified proteoglycans, in tendon fibroblasts cultured in 5, 10, and 50 kPa NorHA hydrogels for 4 days (N = 5 independent samples per group). Variance-stabilizing transformation (VST) expression values were standardized by gene (row z-score) and hierarchically clustered.

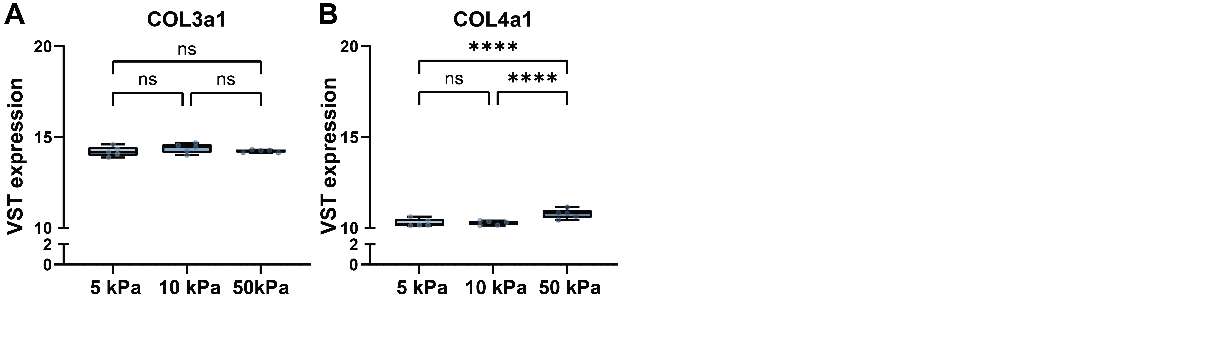

**Supplementary Figure 7. Expression of *Col3a1* and *Col4a1* in 3D tendon fibroblast culture.** VST expression of *Col3a1* and *Col4a1* in tendon fibroblasts cultured in 5, 10, and 50 kPa NorHA hydrogels for 4 days (N = 5 independent samples per group). ****p < 0.0001; ns = not significant; Benjamini-Hochberg-adjusted p values from DESeq2 pairwise differential-expression analysis.
